# CUBE: Multimodal Representation Learning Reveals Biological Structure Across Histomorphology, Spatial Protein Phenotypes, and Transcriptome-Associated Signals

**DOI:** 10.64898/2026.09.14.751379

**Authors:** Zhiyan Ge, Haoyang Cai

## Abstract

Integrating histological, spatial protein, and transcriptomic information into a biologically grounded representation remains challenging because these modalities are rarely available as fully paired measurements, while existing computational approaches are commonly developed around individual modality pairs. To address this, we present CUBE (Colorectal Universal Representation & Bridge Encoder), a multimodal representation-learning framework that uses hematoxylin and eosin (H&E) histology as a bridge to integrate spatial protein phenotypes with transcriptome-associated information from incompletely paired data. CUBE independently learns representations from H&E–multiplex immunohistochemistry (mIHC) and H&E–pseudo-ST relationships and integrates them through attention-based fusion with biological grounding from mIHC-derived concepts. The H&E–mIHC representation supported competitive spatial protein reconstruction, while the pseudo-ST-supervised representation transferred to experimentally measured Visium HD spatial transcriptomics data and improved further after decoder-only calibration with the encoder frozen. Importantly, the fused representation retained biological information beyond its direct training targets, capturing immune–epithelial spatial organization and an independently measured ECM–receptor interaction transcriptomic program. Together, these findings demonstrate that separately paired spatial modalities can be organized through histology into a biologically structured and testable multimodal representation, providing a proof-of-concept strategy for multimodal tissue learning without requiring fully paired molecular measurements.

## 1 Introduction

Histopathology, spatial protein imaging, and spatial transcriptomics provide complementary views of tissue biology. Hematoxylin and eosin (H&E) staining captures tissue architecture, cellular morphology, and spatial organization, whereas multiplex immunohistochemistry (mIHC) provides spatially resolved information on protein phenotypes and cellular components within the tissue microenvironment [1,2]. Spatial transcriptomics (ST) adds a molecular dimension by measuring gene expression with spatial resolution, preserving its localization and organization within tissue [3].Together, these modalities describe tissue state from morphological, protein, and transcriptional perspectives. Although they are not directly interchangeable, information at these levels is biologically coupled, creating an opportunity to learn cross-modal associations and to construct representations that integrate tissue morphology with spatial molecular information [4,5].

Deep learning has enabled increasingly accurate prediction of molecular and phenotypic features from histological images, yet predictive performance alone does not reveal what biological information is encoded in the learned representation [6]. Intermediate features in deep neural networks are typically high-dimensional and difficult to interpret, making it challenging to determine whether a model relies on biologically meaningful tissue signals or on dataset-specific shortcuts [7,8]. In biomedical applications, this distinction is particularly important because a representation becomes more informative when its internal structure can be related to measurable biological properties. Linking latent features to interpretable biological concepts can therefore provide biological grounding for model predictions, help identify the tissue signals associated with those predictions, and provide a more principled basis for model refinement and the generation of testable hypotheses [9,10].

Recent studies have demonstrated that routine H&E histology can be computationally linked to spatial molecular measurements. For spatial protein imaging, HEMIT introduced a cellular-level aligned H&E–mIHC dataset and a translation framework for DAPI, CD3, and panCK [1]. More recent studies have substantially expanded this direction: GigaTIME generates virtual multiplex immunofluorescence profiles spanning 21 protein channels from H&E images [11], while HEX predicts high-dimensional spatial proteomic profiles and further uses the inferred protein information for biological interpretation and biomarker discovery [12]. In parallel, prediction of spatial gene expression from histology has developed from early approaches such as ST-Net [5] to models that explicitly incorporate spatial relationships, including Hist2ST [13]. More recent methods have also introduced multimodal representation learning; for example, mclSTExp uses contrastive learning to align histological and spatial transcriptomic features within a multimodal embedding space [14]. HiST further demonstrated high-fidelity spatial gene-expression reconstruction across multiple cancer types and its utility for downstream biological and clinical analyses [15]. A recent systematic benchmark of eleven H&E-to-ST methods further illustrates the rapid expansion of this field [16]. Despite these advances, existing approaches remain largely organized around individual modality pairs, either linking morphology with spatial protein measurements or linking morphology with transcriptomic profiles. Even when representation learning or biological interpretation is incorporated, it is generally developed within a single cross-modal relationship [12,14,15]. Whether protein-phenotypic and transcriptome-associated information learned from separately paired datasets can be integrated through a common histological bridge into a biologically testable representation remains less explored.

To address this gap, we developed CUBE (Colorectal Universal Representation & Bridge Encoder), a multimodal representation-learning framework that uses H&E histology as a bridge to integrate spatial protein phenotypes and transcriptome-associated information from incompletely paired data. CUBE first learns two H&E-derived representations independently: UR1 from the relationship between H&E and mIHC, and UR2 from the relationship between H&E and DeepSpot-derived pseudo-ST [17]. These representations are subsequently integrated through an attention-based fusion module and biologically grounded using measurable mIHC-derived concepts. In this way, cross-modal prediction serves not only as an end task but also as supervision for learning representations that retain information associated with distinct biological levels. We further evaluate whether the fused representation contains biologically structured information beyond its direct training targets by examining latent-space organization and its ability to decode an independently defined immune–epithelial spatial organization pattern and experimentally measured transcriptomic programs. CUBE therefore provides a proof-of-concept framework for integrating separately paired multimodal tissue information into a biologically testable representation and for evaluating what biological information is retained after cross-modal learning and fusion.

## 2 Results

### 2.1 Overall Architecture of CUBE

CUBE was developed as a multimodal representation-learning framework that uses H&E histology as a bridge modality to connect spatial protein phenotypes with transcriptome-associated information (Figure 1). The framework consists of three principal branches: (i) the H&E–mIHC branch, (ii) the H&E–ST branch, and (iii) the universal representation (UR) fusion/concept prediction branch. The first two branches learn modality-aligned representations from H&E–mIHC relationships and H&E with DeepSpot-derived pseudo-ST supervision, respectively, whereas the third branch integrates the two H&E-derived representations and uses biologically interpretable mIHC-derived concepts to guide the fused representation. During preprocessing, the original 1024 × 1024 H&E and mIHC patches were resized to 512 × 512 for the H&E–mIHC branch, while H&E patches were additionally resized to 256 × 256 for the H&E–ST branch. For each original 1024 × 1024 H&E patch, a 16 × 16 pseudo-ST matrix spanning the top 256 highly variable predicted genes was generated using a pretrained DeepSpot model. For each original 1024 × 1024 mIHC patch, channel-specific Otsu thresholds were independently estimated for DAPI, CD3, and panCK, and the corresponding positive-pixel fractions were used as three continuous concept scores [18].

**Figure 1.**
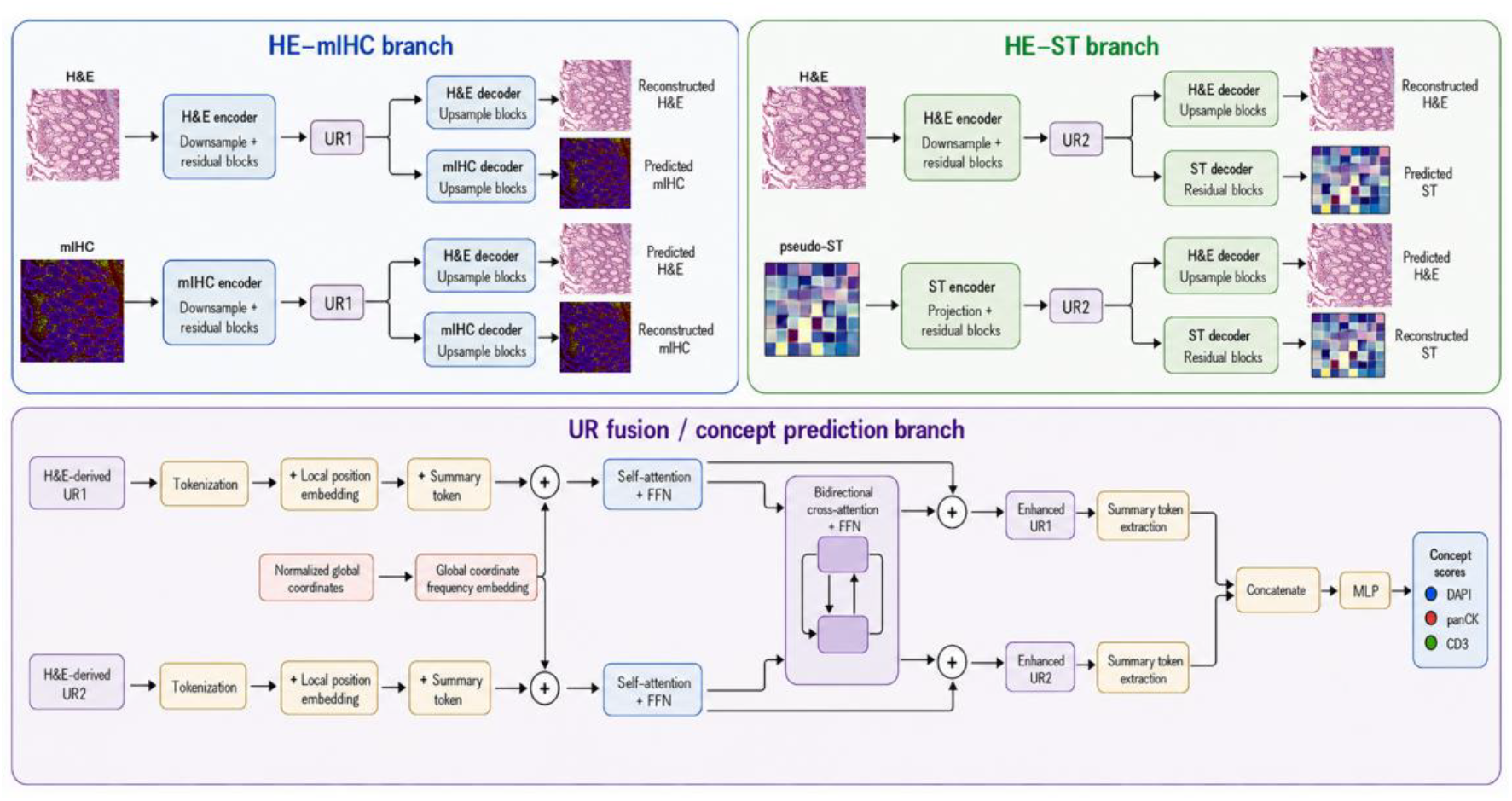
Overview of the CUBE framework. CUBE comprises the H&E–mIHC branch, H&E– ST branch, and universal representation fusion/concept prediction branch. All encoder and decoder instances are independently parameterized; the H&E-to-mIHC pathway uses marker-specific decoders for DAPI, CD3, and panCK, and representation fusion uses only H&E-derived UR1 and UR2. H&E and mIHC panels are real study images manually embedded into the schematic, with gamma adjustment applied to mIHC for visualization. ST matrices are illustrative representations rather than experimental examples. Schematic elements were generated using OpenAI ChatGPT Images 2.0 in ChatGPT Work/Codex and assembled by the authors; biological images were not generatively modified.

The H&E–mIHC branch uses independently parameterized residual convolutional encoders to map H&E and mIHC inputs into H&E-derived and mIHC-derived UR1 representations, respectively. A weak alignment constraint is applied to spatially pooled summaries of the two UR1 representations, encouraging them to retain shared tissue-level information without enforcing strict element-wise correspondence between the complete latent feature maps. Each UR1 representation is subsequently decoded into both modalities through independently parameterized decoders, forming H&E self-reconstruction, H&E-to-mIHC prediction, mIHC-to-H&E prediction, and mIHC self-reconstruction pathways. To accommodate the heterogeneous spatial distributions of the three mIHC markers, the H&E-to-mIHC pathway employs separate marker-specific decoders for DAPI, CD3, and panCK.

The H&E–ST branch follows the same encoder–representation–decoder principle to learn UR2. H&E and pseudo-ST inputs are processed by independent encoders to generate H&E-derived and pseudo-ST-derived UR2 representations, with a weak pooled alignment constraint promoting shared tissue-level information while preserving the spatial structure of each latent representation. The H&E-derived UR2 is decoded for H&E self-reconstruction and H&E-to-pseudo-ST prediction, whereas the pseudo-ST-derived UR2 is decoded for pseudo-ST self-reconstruction and auxiliary pseudo-ST-to-H&E reconstruction. The H&E-to-pseudo-ST pathway provides the primary cross-modal supervision, while the reverse pseudo-ST-to-H&E pathway serves as a lower-weight auxiliary constraint on representation learning.

For representation fusion, H&E-derived UR1 and UR2 extracted from the same tissue patch are used as the two input streams. Each feature map is first tokenized and supplemented with a branch-specific local positional embedding, after which a learnable summary token is prepended. The normalized global coordinates of each tissue patch are encoded using Fourier frequency features and projected into the latent feature space before being added to both token sequences. UR1 and UR2 are then independently refined through self-attention and feed-forward networks, followed by bidirectional cross-attention that enables each representation to incorporate information from the other. The updated summary tokens from the two streams are concatenated to form the final fused UR, which is passed to an MLP-based concept prediction head to estimate the DAPI-, CD3-, and panCK-positive fractions. This concept-based supervision encourages the fused representation to retain nuclear-, immune-, and epithelial-associated phenotypic information in addition to the complementary cross-modal information captured by UR1 and UR2.

### 2.2 CUBE Achieves Accurate H&E-to-mIHC Prediction through a Modality-Aligned Representation

To evaluate whether UR1 retained spatial protein phenotype information that could be recovered from histomorphology, we first assessed H&E-to-mIHC prediction on the 945 held-out test patches from the HEMIT dataset. CUBE was compared with U-Net, ResNet, pix2pix U-Net, pix2pix ResNet, and HEMIT-512 adapted under the same 512 × 512 preprocessing and evaluation pipeline (Figure 2a) [19,20,21,1]. CUBE achieved a mean Pearson correlation of 0.7256, outperforming U-Net (0.5302), ResNet (0.5562), pix2pix U-Net (0.5446), and pix2pix ResNet (0.7084). Its performance was closely comparable to HEMIT-512 adapted, which achieved the highest mean correlation of 0.7310. Thus, although CUBE was not specifically designed as a dedicated virtual-staining model, its H&E–mIHC branch achieved competitive reconstruction performance relative to the task-specific HEMIT-512 adapted benchmark.

**Figure 2.**
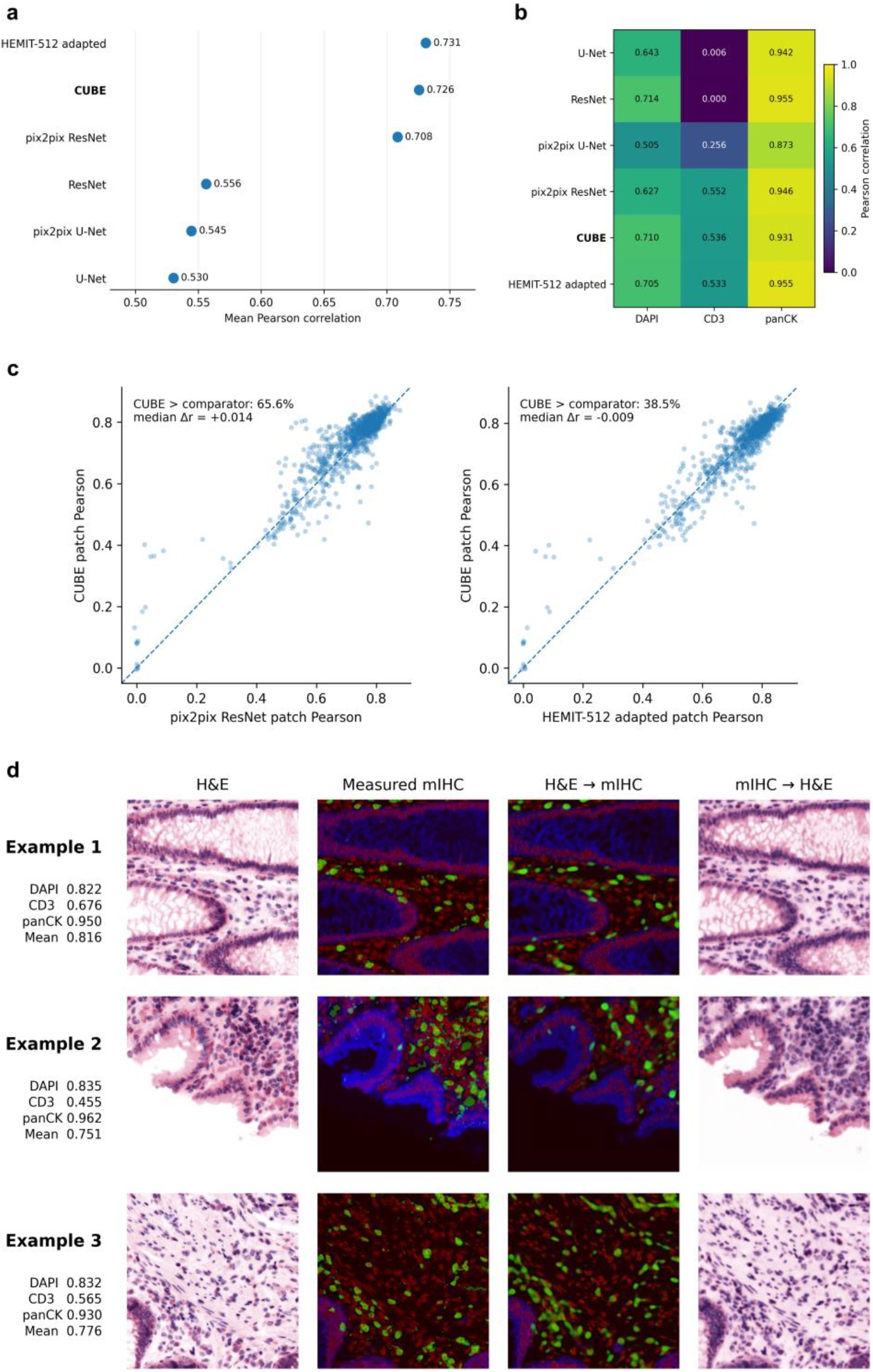
Performance of CUBE in H&E–mIHC cross-modal reconstruction. (a) Mean Pearson correlation of H&E-to-mIHC predictions on 945 held-out test patches for CUBE and five comparator models under the standardized 512 × 512 evaluation setting. (b) Marker-resolved Pearson correlations for DAPI, CD3, and panCK. (c) Paired patch-wise comparison of CUBE with pix2pix ResNet and HEMIT-512 adapted; dashed diagonal lines indicate equal performance between models. (d) Representative bidirectional reconstruction examples across distinct tissue morphologies. Columns show H&E, measured mIHC, H&E-to-mIHC prediction, and mIHC-to-H&E reconstruction. Composite mIHC images display DAPI in blue, CD3 in green, and panCK in red; marker-specific Pearson correlations shown for each example correspond to the H&E-to-mIHC prediction.

Marker-resolved evaluation further showed that CUBE maintained substantial predictive performance across DAPI, CD3, and panCK (Figure 2b). CUBE achieved Pearson correlations of 0.7105 for DAPI, 0.5356 for CD3, and 0.9308 for panCK. Its DAPI performance was close to the best-performing ResNet baseline (0.7141), while its CD3 prediction substantially exceeded U-Net (0.0057) and ResNet (0.0000) and remained comparable to pix2pix ResNet (0.5523) and HEMIT-512 adapted (0.5328). Although the panCK correlation of CUBE was modestly lower than several comparators, it remained high at 0.9308. These results indicate that CUBE maintained strong performance across markers while avoiding the severe degradation observed for the sparse CD3 channel in standard CNN baselines.

We next examined whether CUBE’s higher mean Pearson correlation than pix2pix ResNet was consistently observed across individual test patches (Figure 2c). Compared with pix2pix ResNet, CUBE achieved a higher patch-wise Pearson correlation in 65.6% of test patches, with a median difference of Δr = +0.014. This indicates that the improvement in mean performance was not driven solely by a small subset of exceptionally well-predicted patches. In comparison with HEMIT-512 adapted, CUBE achieved higher patch-wise correlations in 38.5% of test patches, with a median Δr of ™0.009. The relatively small median difference and the close distribution of paired patch-level performances were consistent with the small difference in mean Pearson correlation between CUBE and HEMIT-512 adapted (0.7256 vs 0.7310).

Representative reconstructions further illustrated the cross-modal information retained by UR1 across morphologically distinct tissue regions, including glandular structures and regions with different epithelial and cellular organization (Figure 2d). H&E-to-mIHC predictions reproduced the major spatial distributions of DAPI, CD3, and panCK across these representative regions, while the reverse mIHC-to-H&E pathway qualitatively recovered the principal histological organization of the corresponding tissue. These bidirectional reconstructions indicate that UR1 retains cross-modal information sufficient to support reconstruction of both histomorphology and spatial protein phenotypes.

### 2.3 CUBE Learns a Transferable Transcriptome-Associated Representation from Pseudo-ST Supervision

We first evaluated whether the H&E–ST branch could reconstruct the transcriptome-associated patterns defined by its DeepSpot-derived pseudo-ST supervision. On the 945 held-out HEMIT test patches, CUBE showed consistently high agreement with the pseudo-ST targets across multiple levels of evaluation (Figure 3a). The global Pearson correlation across all predicted values was 0.802, while the mean gene-wise global correlation reached 0.795. When expression was spatially averaged within each patch and compared across samples, the mean cross-sample abundance correlation increased to 0.889. Importantly, the mean within-patch spatial correlation, calculated independently for each gene within each 16 × 16 spatial grid, remained 0.723. These results indicate that CUBE reproduced not only overall gene abundance differences among tissue patches but also the local spatial organization encoded in the teacher-defined pseudo-ST targets.

**Figure 3.**
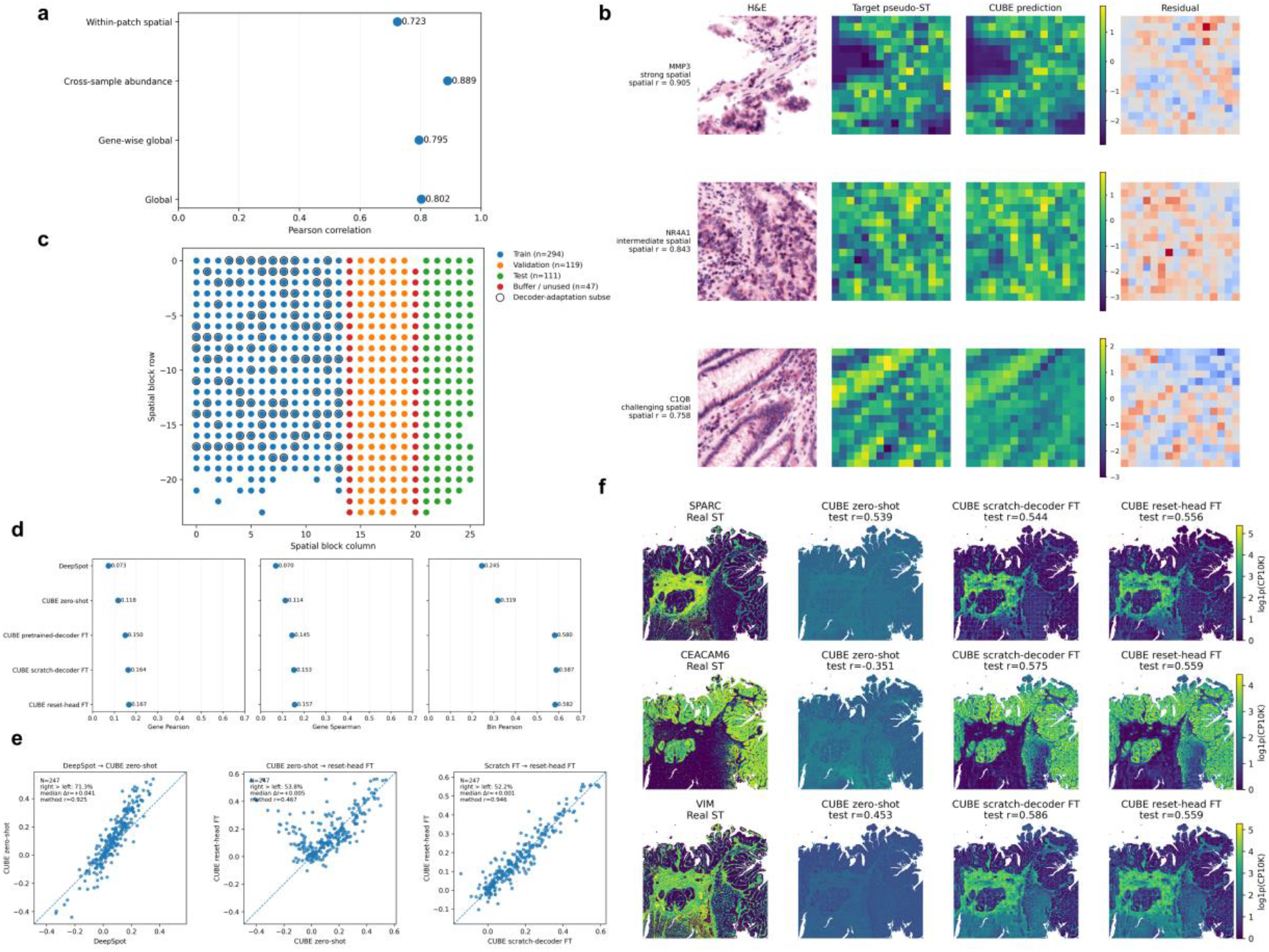
Performance of CUBE in pseudo-ST reconstruction and transfer to measured Visium HD spatial transcriptomics. a) Pearson correlation performance of H&E-to-pseudo-ST reconstruction on 945 held-out HEMIT test patches, evaluated by global, gene-wise global, cross-sample abundance, and within-patch spatial correlations.b)Representative H&E-to-pseudo-ST spatial reconstructions for MMP3, NR4A1, and C1QB. Columns show the H&E patch, DeepSpot-derived target pseudo-ST, CUBE prediction, and residual map. Spatial *r* denotes the within-patch Pearson correlation between the target and predicted expression fields. c) Contiguous spatial holdout design for the Visium HD section, including training (n = 294), validation (n = 119), test (n = 111), and unused buffer (n = 47) regions. Circled points indicate the 100-patch decoder-adaptation subset sampled from the training region. During real-ST adaptation, the H&E-to-UR2 encoder was frozen and only the ST decoder was updated. d) Performance on the matched Visium HD held-out test region for DeepSpot, CUBE zero-shot, pretrained-decoder fine-tuning, scratch-decoder fine-tuning, and reset-head fine-tuning, evaluated by gene-wise Pearson correlation, gene-wise Spearman correlation, and bin-wise Pearson correlation. e) Paired per-gene Pearson correlation comparisons on 247 evaluable genes between DeepSpot and CUBE zero-shot, CUBE zero-shot and reset-head fine-tuning, and scratch-decoder and reset-head fine-tuning. Dashed diagonal lines indicate equal performance between paired methods. f) Whole-slide qualitative expression maps for SPARC, CEACAM6, and VIM, showing measured Visium HD expression, CUBE zero-shot prediction, scratch-decoder fine-tuning, and reset-head fine-tuning. Color bars indicate log1p(CP10K) expression values. Reported *r* values were calculated exclusively on the held-out test region.

Representative examples further illustrated this spatial reconstruction capability (Figure 3b). Across genes with different levels of reconstruction difficulty, including MMP3, NR4A1, and C1QB, the predicted 16 × 16 expression fields preserved major spatial patterns observed in the corresponding pseudo-ST targets, with within-patch spatial correlations of 0.905, 0.843, and 0.758, respectively. The residual maps further visualized local differences between the predicted and target expression fields. Together, these quantitative and qualitative results confirmed that the H&E-derived UR2 effectively captured the pseudo-transcriptomic structure provided by DeepSpot supervision.

Because pseudo-ST is itself inferred from H&E rather than independently measured, we next asked whether the learned UR2 contained information transferable to experimentally measured spatial transcriptomics. We therefore evaluated CUBE on the 10x Genomics colorectal cancer Visium HD section using a contiguous spatial holdout design (Figure 3c) [22,23]. The section was divided into spatially separated training, validation, and test regions with unused buffer columns between them, yielding 294 training-region patches, 119 validation patches, 111 held-out test patches, and 47 buffer patches. A subset of 100 patches from the training region was used for real-ST decoder adaptation. Throughout all adaptation experiments, the pretrained H&E-to-UR2 encoder was kept completely frozen, such that only the mapping from the fixed UR2 representation to measured ST was recalibrated. This design allowed us to assess whether limited zero-shot performance primarily reflected a mismatch between the pseudo-ST-trained decoder and the measured-ST domain rather than an absence of the transferable information in UR2.

On the same 111-patch held-out spatial test region, zero-shot CUBE outperformed its DeepSpot teacher across all three evaluation metrics (Figure 3d). DeepSpot achieved gene-wise Pearson and Spearman correlations of 0.0730 and 0.0702, respectively, with a bin-wise Pearson correlation of 0.2447. Without exposure to measured ST during training, CUBE zero-shot increased these values to 0.1177, 0.1140, and 0.3189, respectively. Decoder adaptation using only the 100 real-ST training patches further improved performance while leaving the H&E-to-UR2 encoder unchanged. Fine-tuning the pretrained decoder increased the gene-wise Pearson correlation to 0.1503, whereas training a newly initialized decoder from the frozen UR2 reached 0.1641. Resetting the final output layer of the pretrained decoder before fine-tuning produced a similar value of 0.1670. The corresponding gene-wise Spearman correlations were 0.1447, 0.1535, and 0.1566, and bin-wise Pearson correlations were 0.5801, 0.5873, and 0.5820. These improvements indicate that the frozen UR2 retained information that could be decoded into measured ST after limited real-ST calibration. The higher performance of scratch-decoder and reset-head fine-tuning relative to direct fine-tuning of the pretrained decoder further suggests that the pseudo-ST-trained decoder mapping contributed to the domain mismatch, while the similar performance of scratch-decoder and reset-head adaptation indicates no clear additional advantage from retaining the pretrained decoder trunk.

Gene-level paired comparisons showed that the zero-shot improvement over DeepSpot was broadly distributed rather than being driven by a small number of genes (Figure 3e). Among 247 genes eligible for paired evaluation, CUBE zero-shot achieved a higher Pearson correlation than DeepSpot for 71.3% of genes, with a median improvement of Δr = +0.041. Moreover, gene-wise performances of the two methods remained strongly correlated (r = 0.925), indicating that CUBE retained much of the teacher-defined relative structure of easier and more difficult genes while shifting performance upward on the measured-ST test region. Real-ST adaptation produced additional, but more gene-dependent, changes: reset-head fine-tuning improved over zero-shot CUBE for 53.8% of genes with a median Δr of +0.005, whereas reset-head and scratch-decoder fine-tuning were nearly indistinguishable, with reset-head outperforming scratch for 52.2% of genes and a median Δr of only +0.001. These results support transfer beyond direct reproduction of the pseudo-ST teacher signal while also showing that measured-ST adaptation does not uniformly benefit every gene.

Whole-slide qualitative maps provided further insight into the nature of this pseudo-to-real domain gap (Figure 3f). Zero-shot predictions could retain the coarse spatial geometry of measured gene-expression patterns, but their expression contrast was compressed and, for some genes, partially inverted. For example, the zero-shot maps of SPARC and VIM preserved major spatial organization despite reduced dynamic range, whereas CEACAM6 showed an inverted expression relationship within the held-out test region. After decoder adaptation, gene-specific contrast and spatial correspondence to measured ST were substantially restored, as illustrated by the marked improvement of CEACAM6 from a zero-shot test correlation of −0.351 to 0.575 and 0.559 after scratch-decoder and reset-head fine-tuning, respectively. Similar improvements were observed for SPARC and VIM. The whole-slide maps are shown only for qualitative visualization, whereas all reported correlations were calculated exclusively on the untouched 111-patch spatial test region. Collectively, these results indicate that pseudo-ST supervision enabled CUBE to learn a transcriptome-associated UR2 that retained transferable spatial information, while improved mapping to measured Visium HD expression required calibration of the downstream decoder to the target ST domain.

### 2.4 CUBE Integrates Complementary UR1 and UR2 Representations for Phenotypic Concept Prediction

We next examined whether UR1 and UR2 provided complementary information for predicting the mIHC-derived phenotypic concept scores. To assess the contribution of representation fusion, we performed an ablation analysis using independently trained, architecture-matched UR1-only and UR2-only models. On the validation set, Fusion achieved an overall concept MSE of 0.0066, representing reductions of 25.1% and 24.4% relative to UR1-only (0.0088) and UR2-only (0.0087), respectively (Figure 4a). Fusion also achieved the highest Pearson correlation for each concept, reaching 0.452 for DAPI, 0.883 for CD3, and 0.936 for panCK, compared with 0.318, 0.879, and 0.922 for UR1-only and 0.369, 0.766, and 0.910 for UR2-only. Notably, UR2-only outperformed UR1-only for DAPI despite the concept labels being derived directly from mIHC. These results indicate that information relevant to the mIHC-derived phenotypic concepts was not confined to the mIHC-supervised UR1, and that the transcriptome-associated UR2 provided complementary information that could be exploited through representation fusion.

**Figure 4.**
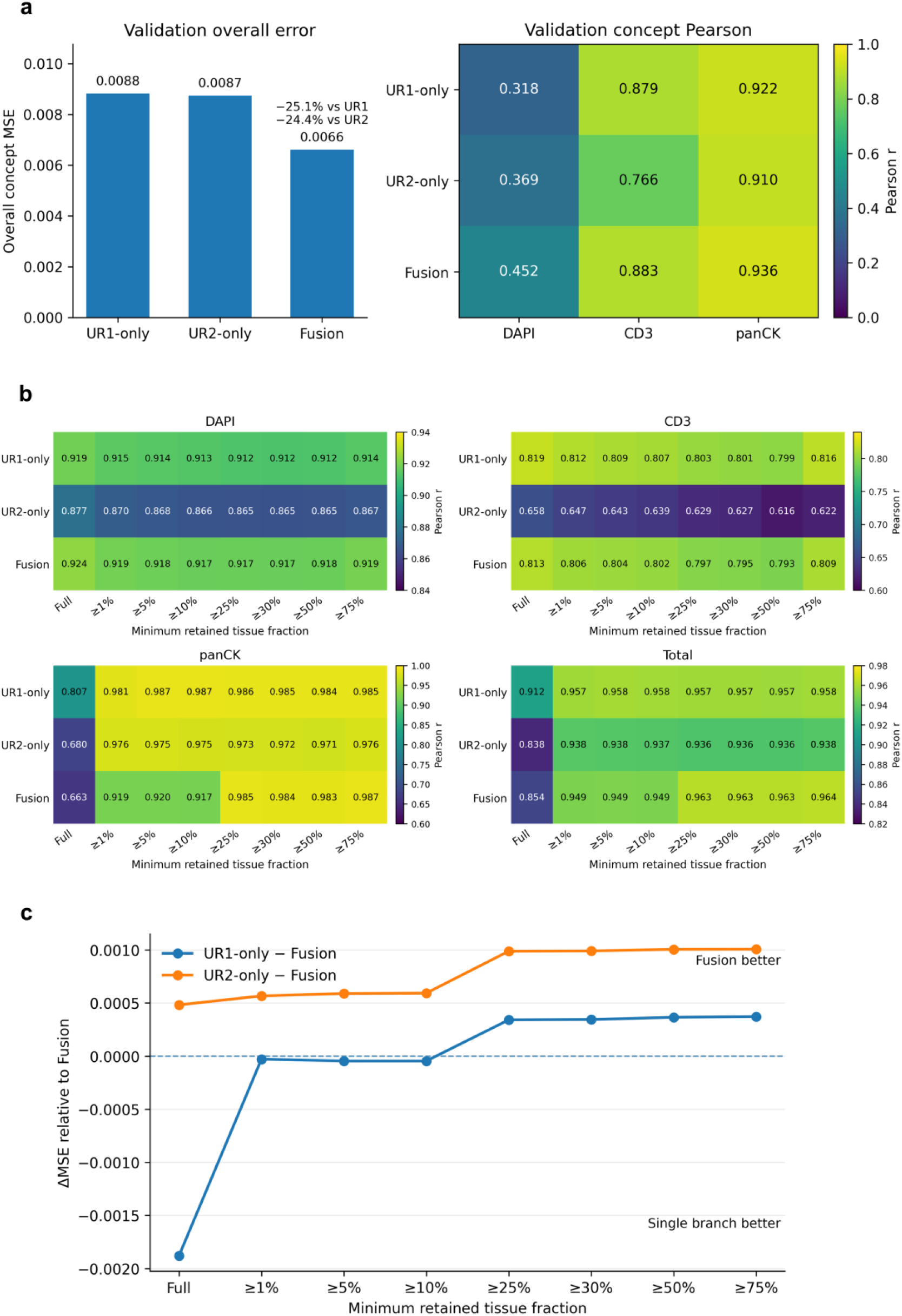
Complementary UR1 and UR2 representations improve phenotypic concept prediction. a) Validation performance of UR1-only, UR2-only, and Fusion models for prediction of mIHC-derived concept scores. Left, overall concept MSE; right, Pearson correlations for DAPI, CD3, and panCK. b) Held-out test Pearson correlations for DAPI, CD3, panCK, and total concept prediction across progressively stricter minimum retained tissue-fraction thresholds. c) Difference in MSE between each single-representation model and Fusion across the same tissue-content thresholds.

When the models were subsequently evaluated on the full held-out test set, however, the pattern differed from that observed on the validation set: Fusion showed a lower total Pearson correlation than UR1-only (0.854 versus 0.912), although it remained above UR2-only (0.838). This discrepancy prompted us to examine whether variation in retained tissue content contributed to the change in relative performance. We therefore re-evaluated all three models across progressively stricter tissue-content thresholds (Figure 4b). As low-tissue patches were progressively excluded, the relative performance of Fusion improved. Fusion consistently achieved slightly higher DAPI correlations than UR1-only, whereas UR1-only remained modestly stronger for CD3. The strongest tissue-content dependence was observed for panCK, for which Fusion performance increased markedly after low-tissue patches were excluded. Consequently, the total Pearson correlation of Fusion surpassed UR1-only from the ≥25% threshold onward, reaching 0.963 compared with 0.957 for UR1-only and 0.936 for UR2-only at this threshold.

The same tissue-dependent pattern was observed when performance was assessed using MSE differences relative to Fusion (Figure 4c). The difference between UR2-only and Fusion remained positive across all tissue-content thresholds, indicating consistently lower MSE for Fusion than for UR2-only. In contrast, the UR1-only–Fusion difference was negative under the full-set and lower tissue-content conditions, approached zero as low-tissue samples were progressively removed, and became positive from the ≥25% threshold onward. The fusion advantage over UR1 then remained stable across increasingly tissue-enriched subsets. Together, these results show that UR1 and UR2 provide complementary information for phenotypic concept prediction, while the relative benefit of fusion depends on the amount of tissue content retained in the input patch.

Positive ΔMSE values indicate lower error for Fusion, whereas negative values indicate lower error for the corresponding single-representation model; the dashed horizontal line denotes equal MSE.

### 2.5 The Fused CUBE Representation Encodes Spatial Phenotypic Structure and Retains Transcriptome-Associated Information

We next investigated whether the fused CUBE representation retained biologically structured information beyond that required for direct reconstruction and concept prediction. Because global spatial coordinates were explicitly incorporated during representation fusion, we first generated fixed-coordinate fused representations by replacing the coordinates of all 5292 HEMIT patches with the same dataset-level mean coordinate while keeping the trained fusion model unchanged. The resulting 512-dimensional representations were reduced by PCA and visualized using UMAP (Figure 5a) [24,25]. The latent geometry exhibited nonuniform distributions of DAPI, CD3, panCK, and tissue content, with continuous phenotypic gradients and locally enriched regions distributed across the embedding. Partial split-associated structure was also visible, consistent with residual dataset-level heterogeneity. Because DAPI-, CD3-, and panCK-positive fractions had been used as supervision during fusion training, these associations were interpreted descriptively rather than as independent biological validation. To further characterize the organization of this representation space, we applied K-means clustering to the PCA-reduced fixed-coordinate representations and used an eight-cluster partition as a higher-resolution exploratory representation of latent tissue states (Figure 5b) [26].

**Figure 5.**
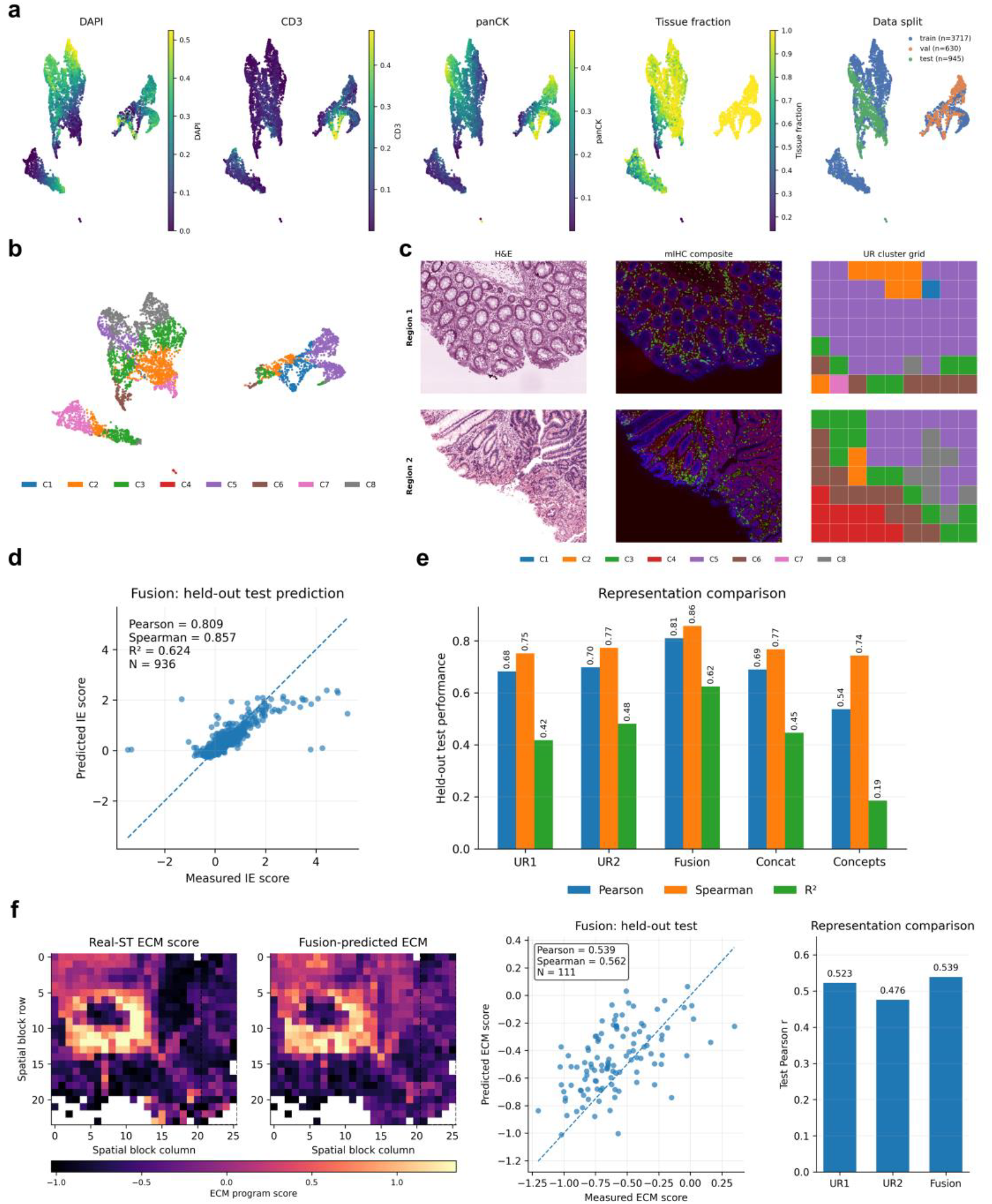
Biological structure and downstream information encoded by the fused CUBE representation. (a) Fixed-coordinate UMAP visualization of fused UR representations from all 5292 HEMIT patches. Global spatial coordinates were replaced by the same dataset-level mean coordinate before representation extraction. Points are colored by DAPI-, CD3-, and panCK-positive fractions, retained tissue fraction, and data split. (b) Eight-cluster K-means partition of the PCA-reduced fixed-coordinate fused representations shown in (a). Colors denote cluster assignments C1–C8. (c) Spatial mapping of fused-UR cluster assignments in two representative registered tissue regions. Columns show the corresponding H&E image, mIHC composite, and patch-level UR cluster grid. Cluster colors correspond to those in (b); each grid cell represents one tissue patch rather than a pixel-level segmentation. (d) Decoding of an immune–epithelial spatial enrichment (IE) score from frozen representations. Left, measured versus Fusion-predicted IE scores on 936 evaluable held-out test patches, with Pearson correlation, Spearman correlation, and R^2^. Right, held-out prediction performance using UR1, UR2, Fusion, simple concatenation of pooled UR1 and UR2 features (Concat), or the three supervised abundance concept scores (Concepts). The IE score quantifies preferential localization of CD3-associated signals within or near panCK-rich epithelial neighborhoods. (e) External real-ST validation of ECM-associated transcriptomic information using a colorectal cancer Visium HD section. Left, spatial maps of the experimentally measured and Fusion-predicted 17-gene KEGG ECM–receptor interaction (hsa04512) program score. Middle, measured versus Fusion-predicted ECM scores on the 111 held-out test patches. Right, test Pearson correlations obtained from frozen UR1, UR2, and Fusion representations.

We then examined whether these unsupervised representation states corresponded to spatially organized tissue patterns by mapping cluster assignments back to their registered tissue regions (Figure 5c). Each cluster label represented the fused UR of an individual patch rather than a pixel-level segmentation. Across representative regions, cluster assignments formed spatially coherent patch-level domains that qualitatively corresponded to differences in glandular architecture, epithelial organization, cellularity, and tissue absence observed in the paired H&E and mIHC images. Different tissue patterns therefore showed structured localization within the fused representation space, and the corresponding latent states could be mapped back to spatially coherent regions within registered tissue samples. These observations suggest that the fused representation organized histomorphological and protein-phenotypic variation into spatially meaningful representation states, although the clusters were not treated as formally annotated pathological classes.

To test whether the fused representation retained spatial protein information that had not been directly used as a fusion target, we next defined an immune–epithelial spatial enrichment (IE) score (Figure 5d). Unlike the DAPI, CD3, and panCK abundance scores used during fusion training, the IE score quantified the spatial relationship between CD3- and panCK-associated signals. Specifically, for each CD3-positive pixel, the local fraction of panCK-positive pixels within a 20 μm circular neighborhood was calculated, averaged across all CD3-positive pixels, normalized by the global panCK-positive fraction of the patch, and log2-transformed. Higher IE scores therefore indicate preferential localization of CD3-associated signals within or near panCK-rich epithelial neighborhoods. The score was designed as a quantitative surrogate for an immune–epithelial infiltration- or interaction-associated spatial phenotype rather than as a direct measurement of immune infiltration. Using the frozen representations as inputs to lightweight prediction probes, Fusion predicted the held-out IE score with a Pearson correlation of 0.809, a Spearman correlation of 0.857, and an R^2^ of 0.624 across 936 evaluable test patches. Its Pearson correlation exceeded those obtained from UR1 alone (0.683), UR2 alone (0.698), a simple concatenation of pooled UR1 and UR2 features (0.689), and a predictor using only the three supervised abundance concept scores (0.537). These results indicate that Fusion retained immune–epithelial spatial information that could not be explained by marker abundance alone.

Finally, we asked whether the fused representation also retained transcriptome-associated biological information measured independently from the pseudo-ST supervision used during CUBE training. We evaluated frozen CUBE representations on an external colorectal cancer Visium HD section using a 17-gene KEGG ECM–receptor interaction program derived from experimentally measured spatial transcriptomics (Figure 5e) [27]. A patch-level ECM program score was constructed from the matched pathway genes, and representation probes were trained on the spatial training region and evaluated on the 111 held-out test patches. The spatial distribution predicted from Fusion reproduced major regional variation in the measured ECM program, with a test Pearson correlation of 0.539 and Spearman correlation of 0.562. Fusion also achieved the highest Pearson correlation among the three CUBE representations, compared with 0.523 for UR1 and 0.476 for UR2. Importantly, the ability to decode an independently measured pathway-level transcriptomic signal from the frozen fused representation provides evidence that CUBE retained transcriptome-associated information that extended beyond the mIHC-derived concept supervision. Together, the latent-space geometry, spatial clustering, immune–epithelial enrichment decoding, and external real-ST analysis indicate that the fused CUBE representation integrates biologically structured information spanning histomorphology, spatial protein phenotypes, and transcriptome-associated programs.

## 3 Discussion

Tumor tissue state is reflected across multiple, related but non-equivalent biological levels, including transcriptional programs, spatial protein phenotypes, and histomorphology. Learning the relationships among these levels is valuable for constructing representations that capture tissue biology more broadly than any single measurement modality. In practice, however, sufficiently large datasets in which H&E, multiplex protein imaging, and spatial transcriptomics are all measured in a fully paired manner remain difficult to obtain [4,1]. CUBE was developed for this incompletely paired setting, using H&E as a common bridge between H&E–mIHC and H&E– pseudo-ST relationships. The framework asks whether protein-phenotypic and transcriptome-associated information can first be learned separately from these paired relationships and then integrated into a biologically decodable representation. Across the present experiments, UR1 retained recoverable spatial protein information, UR2 retained molecular information transferable to experimentally measured spatial transcriptomics, and Fusion preserved complementary biological signals beyond the quantities directly used for supervision. In CUBE, the “universal” representation therefore refers to the integration of histomorphological, protein-phenotypic, and transcriptome-associated information within a common representation framework.

The use of separate UR1 and UR2 representations follows directly from the structure of the available supervision. H&E–mIHC and H&E–pseudo-ST provide two distinct cross-modal learning signals, so CUBE first learns branch-specific representations and integrates the resulting H&E-derived features only at a higher level. This staged formulation preserves information emphasized by each cross-modal relationship before fusion. The H&E–mIHC results provide evidence that UR1 retains meaningful protein-phenotypic information: although CUBE was not designed as a dedicated virtual-staining model, it achieved a mean H&E-to-mIHC Pearson correlation of 0.7256, closely approaching the 0.7310 obtained by the adapted task-specific HEMIT model (Figure 2a). Marker-resolved results further showed that this information was recoverable across DAPI, CD3, and panCK. These results support UR1 as a biologically informative representation within the broader multimodal framework.

The role of pseudo-ST supervision requires a different interpretation. Because the pseudo-ST targets were inferred from H&E by DeepSpot, high agreement with these targets alone cannot establish transfer to experimentally measured transcriptomic information. The Visium HD experiments therefore provide the more informative test of UR2. On the same spatially held-out test region, DeepSpot achieved a gene-wise Pearson correlation of 0.073, whereas CUBE reached 0.118 without exposure to measured ST during training (Figure 3d). When the H&E-to-UR2 encoder was kept completely frozen and only the mapping from UR2 to measured ST was recalibrated, performance increased to approximately 0.15–0.17 depending on the decoder adaptation strategy (Figure 3d). The zero-shot improvement was also broadly distributed across genes, with CUBE outperforming DeepSpot for 71.3% of evaluable genes (Figure 3e). Notably, newly initialized and reset-head decoders performed at least as well as direct fine-tuning of the pretrained pseudo-ST decoder. This pattern suggests that pseudo-ST supervision may be particularly useful for shaping a transferable representation, while limited real-ST adaptation can further improve the mapping from this representation to measured expression. Pseudo-ST should therefore be viewed as a source of transcriptome-associated representation supervision or inductive bias, with experimentally measured ST remaining necessary for direct validation and target-domain calibration.

The fusion experiments further indicate that UR1 and UR2 contain complementary information, although the benefit of fusion depends on the amount of informative tissue present in the input. On the validation set, Fusion reduced concept-prediction error compared with both UR1-only and UR2-only models (Figure 4a). On the complete held-out test set, UR1 achieved a higher overall Pearson correlation than Fusion; however, Fusion improved progressively as low-tissue patches were excluded and surpassed UR1 when the analysis was restricted to patches with at least 25% retained tissue (Figure 4b,c). This pattern suggests that representation complementarity becomes most useful when sufficient tissue-derived information is available. The downstream analysis provide additional support for the biological content of the fused representation. Fusion predicted the immune–epithelial spatial enrichment score, which captures the spatial relationship between CD3- and panCK-associated signals rather than their individual abundances, substantially better than a predictor using only the three supervised abundance concepts (Figure 5d). Fusion also retained experimentally measured ECM-associated transcriptomic information in the external Visium HD analysis and achieved the highest Pearson correlation among UR1, UR2, and Fusion (Figure 5e). Together, these results indicate that the fused representation preserves spatial and molecular information that extends beyond direct memorization of the DAPI-, CD3-, and panCK-positive fractions used during training.

Several limitations define the scope of the present study. CUBE was developed primarily using colorectal cancer data, and the mIHC branch included only DAPI, CD3, and panCK, providing a limited view of nuclear-, immune-, and epithelial-associated phenotypes. The corresponding concept scores were defined as marker-positive pixel fractions and should not be interpreted as direct cell counts or cell-type annotations. Likewise, the IE score is a surrogate measure of local immune–epithelial spatial enrichment and does not directly measure immune infiltration or physical cell–cell interaction. The transcriptomic component remains dependent on DeepSpot-derived pseudo-ST, while evaluation against experimentally measured Visium HD data was performed in a limited setting. Broader generalization across cohorts, patients, platforms, and tissue types therefore remains to be established. More fundamentally, morphology-based prediction is expected to be strongest for molecular programs that leave detectable spatial or structural signatures in tissue architecture, whereas biological variation with weak morphological correlates may remain difficult to recover from H&E alone [16,6].

Future studies can test the generality of this bridge-modality formulation using richer multiplex protein panels, independent real-ST cohorts and platforms, and larger collections spanning additional cancer and tissue types. Such datasets would enable more rigorous evaluation of whether the observed complementarity generalizes across biological and technical domains and whether more detailed cellular and neighborhood-level phenotypes can be represented. As larger collections of separately paired multimodal data become available, H&E-based bridge learning may provide a scalable strategy for integrating heterogeneous spatial measurements without requiring every modality to be acquired from the same tissue specimen. CUBE therefore serves as a methodological proof-of-concept that separately paired multimodal information can be organized through H&E into complementary representations that remain biologically decodable after fusion.

## 4 Experimental Section

### 4.1 Dataset and Preprocessing

#### (a) HEMIT dataset and data organization

CUBE was primarily developed using the HEMIT colorectal cancer dataset, which contains cellular-level registered H&E and multiplex immunohistochemistry (mIHC) image pairs [1,28]. The dataset comprises 5292 paired 1024 × 1024 image patches, including 3717 training, 630 validation, and 945 held-out test patches. The original HEMIT partition was retained throughout the study without additional reassignment. Spatial coordinates encoded in the patch filenames were parsed for subsequent spatial representation analysis. Global coordinates were normalized using fixed coordinate bases of 22,541 and 64,216 for the two axes, respectively.

#### (b) H&E and mIHC image preprocessing:&

Each 1024 × 1024 H&E patch was resized to 512 × 512 pixels for the H&E–mIHC branch and to 256 × 256 pixels for the H&E–ST branch. The corresponding mIHC patch was resized to 512 × 512 pixels. Area interpolation was used for resizing, and image intensities were scaled from the original 8-bit range to [0,1]. No additional intensity normalization was applied during model loading.

#### (c) mIHC-derived concept score calculation

Three continuous phenotypic concept scores were calculated from the original 1024 × 1024 mIHC patch images before resizing, corresponding to DAPI-, CD3-, and panCK-associated signals. For each channel cc, an independent Otsu threshold T_c_ was estimated from the corresponding 1024 × 1024 single-channel image by maximizing the between-class variance between pixels below and above the candidate threshold.

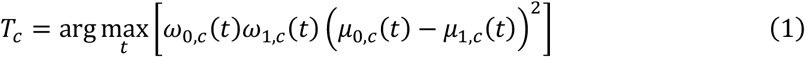

where ω_0,c_(t) and ω_1,c_(t) denote the fractions of pixels assigned to the two intensity classes at candidate threshold tt, and μ_0,c_(t) and μ_1,c_(t) denote their corresponding mean intensities. Pixels with intensities greater than T_c_ were considered marker-positive. The concept score for channel cc was then defined as the proportion of marker-positive pixels within 1024 × 1024 patch,

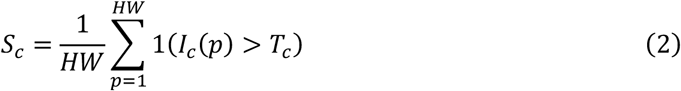

where H=W=1024, I_c_(p) is the intensity of pixel p in channel c, and 1(⋅) is an indicator function that equals 1 when the enclosed condition is satisfied and 0 otherwise. The resulting scores therefore represent marker-positive pixel fractions rather than direct cell counts.

#### (d) DeepSpot-derived pseudo-ST generation

Pseudo-ST targets for the H&E–ST branch were generated from the original 1024 × 1024 H&E patches using pretrained DeepSpot [17]. Inference used the Colon_HEST1K expression model together with the UNI morphology feature extractor [29,30]. A regular 16 × 16 prediction grid was constructed for each patch, yielding 256 spatial positions with a center-to-center spacing of 64 pixels. DeepSpot inference used nine mini-tiles per prediction position, a neighborhood radius of one, and disabled super-resolution inference.

Genes designated as predictable by the pretrained DeepSpot model were ranked according to highly_variable_rank, and the top 256 genes were retained in a fixed order. Predictions were consequently represented as 16 × 16 × 256 pseudo-ST tensors. These targets were inferred from H&E rather than independently measured spatial transcriptomics; experimentally measured ST data were therefore used separately for transfer and external validation analysis.

#### (e) Pseudo-ST normalization

Gene-wise normalization statistics were subsequently calculated using the 3717 HEMIT training patches only. For each of the 256 genes, the mean and population standard deviation were estimated across all 3717×16×16=951,552 training spatial positions. For a pseudo-ST value x_p,g_ at spatial position p and gene g, the normalized value used by the H&E–ST branch was

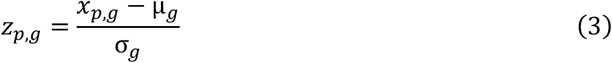

where μ_g_ and σ_g_ were calculated exclusively from the training split. To prevent numerical instability, standard deviations smaller than 10^-6^ were replaced by 1.0 before normalization. The same training-derived statistics and the same fixed 256-gene order were applied to the validation and test sets.

### 4.2 H&E–mIHC Representation Learning

#### (a) Model architecture

The H&E–mIHC branch learned a modality-aligned representation, UR1, from paired H&E and mIHC images. Two structurally identical but independently parameterized encoders received 3 × 512 × 512 H&E and mIHC inputs, respectively. Each encoder contained four stride-2 downsampling blocks with channel dimensions of 32, 64, 128, and 256, followed by two residual blocks. Each downsampling block consisted of a 3 × 3 convolution, Group Normalization, GELU activation, and a residual block [31,32,20]. The resulting H&E- and mIHC-derived representations both had dimensions of 256 × 32 × 32.

Each UR1 representation supported both self- and cross-modal reconstruction, yielding four routes: H&E→H&E, H&E→mIHC, mIHC→H&E, and mIHC→mIHC. The H&E→mIHC route used three independent single-channel decoders for DAPI, CD3, and panCK, whereas the remaining routes used independent three-channel decoders. Each decoder contained four twofold bilinear upsampling blocks with channel dimensions of 256→128→64→32→32, followed by a 3 × 3 output convolution and sigmoid activation. No long-range encoder–decoder skip connections were used, ensuring that reconstruction depended on information encoded in UR1.

#### (b) Training objectives

The H&E–mIHC branch was jointly optimized using self-reconstruction, cross-modal reconstruction, and weak representation-alignment objectives. H&E reconstruction combined pixel-wise L_1_ loss with structural similarity loss [33],

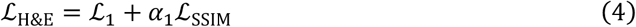

where α_1_ controls the contribution of the structural similarity term.

For mIHC reconstruction, foreground-aware weighted L_1_ loss was used to reduce background dominance. For H&E→mIHC prediction, the per-marker loss ℓ_c_ combined foreground-weighted L_1_, SSIM, multi-scale Pearson correlation, and a CD3-specific soft Dice term [34],

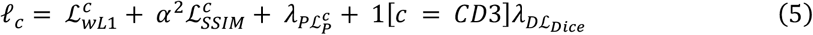

The multi-scale Pearson term integrated correlations evaluated at 512 × 512, 256 × 256, and 128 × 128 resolutions. The corresponding scale weights were 0.5, 0.3, and 0.2. The Pearson-loss weight λ_P_ was set to 0.75, and the CD3-specific Dice weight λ_D_ was set to 0.2. The foreground threshold was 0.05, with an additional foreground weight of 2.0. The SSIM contribution was controlled by α_2_=1.

Marker-specific losses were combined as

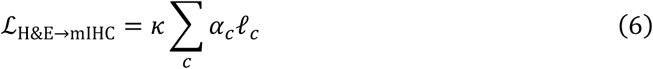

where a_c_ denotes the marker-specific channel weight and κ is a normalization factor used to preserve the overall scale of the three-channel reconstruction loss. Channel weights for DAPI, CD3, and panCK were set to 1, 2, and 1, respectively, with κ=3/Σ_c_a_c_. For mIHC self-reconstruction, the same foreground-aware L_1_, SSIM, and channel-weighting scheme was used, whereas the Pearson and Dice terms were omitted.

To weakly align the H&E- and mIHC-derived UR1 representations without imposing element-wise correspondence, global average pooling was applied over the spatial dimensions and the pooled representations were constrained using mean squared error,

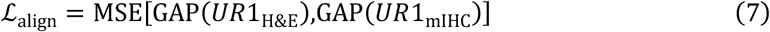

The complete objective was

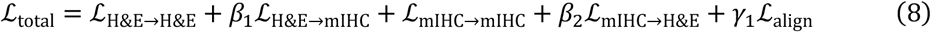

In the final model, β_1_=β_2_=1 and γ_1_=0.05, such that the weak UR1-alignment term remained substantially smaller than the reconstruction objectives.

#### (c) Model training

The branch was trained on 3717 HEMIT training pairs and monitored on 630 validation pairs; the 945 test pairs were excluded from model optimization and checkpoint selection. During training, paired spatial augmentation was applied identically to each H&E–mIHC pair, including random rotations by integer multiples of 90° and independent horizontal and vertical flips with probabilities of 0.5. No spatial augmentation was applied during validation or testing.

Training was performed for 60 epochs using AdamW with a learning rate of 5×10^−5^, weight decay of 1×10^−5^, and a batch size of 4 [35]. At each epoch, H&E→mIHC predictions on the validation set were evaluated separately for DAPI, CD3, and panCK using patch-wise Pearson correlation. The three marker-wise means were averaged. The checkpoint with the highest validation mean Pearson correlation was retained for subsequent evaluation.

### 4.3 H&E–mIHC Benchmarking and Evaluation

#### (a) Evaluation protocol and metrics

H&E→mIHC reconstruction performance was evaluated on the 945 held-out HEMIT test patches. For each patch and each marker channel (DAPI, CD3, and panCK), the predicted and target 512 × 512 images were flattened across spatial pixels and evaluated using Pearson correlation. If either the prediction or target had effectively zero variance (<10^-12^), the corresponding correlation was assigned a value of zero. Marker-level performance was obtained by averaging the per-patch correlations across the test set. An overall score was calculated by first taking the unweighted mean of the DAPI, CD3, and panCK correlations for each patch and then averaging across all test patches.

For paired model comparison, per-patch three-marker mean correlations were additionally calculated for CUBE and selected benchmarks on the same test patches. The difference was defined as Δr=r_CUBE_−r_benchmark_, from which the proportion of patches favoring CUBE and the median Δr were summarized.

#### (b) Benchmark models

CUBE was compared with four standard image-to-image reconstruction baselines: U-Net, ResNet, pix2pix with a U-Net generator, and pix2pix with a ResNet generator. U-Net and ResNet were trained using L_1_ reconstruction loss, whereas the two pix2pix models used a least-squares adversarial objective together with an L_1_ reconstruction term weighted by 30 [36]. All four baselines used the same 512 × 512 H&E–mIHC data and HEMIT train/validation/test partitions as CUBE. Models were trained for 50 epochs using Adam with a learning rate of 3×10^-5^ and batch size 2, and the checkpoint with the highest validation three-marker mean Pearson correlation was retained [37].

We additionally evaluated an adapted implementation of HEMIT, denoted HEMIT-512 adapted. The original HEMIT architecture and training strategy were transferred to the CUBE preprocessing pipeline, with the input resolution reduced from 1024 × 1024 to 512 × 512 and the feature-matching top-k parameter reduced from 1000 to 250 to preserve sampling density after the spatial downscaling. The model used the same HEMIT data partitions and marker ordering as CUBE and was trained for 80 epochs in FP32. Because of these resolution and preprocessing adaptations, this comparator was treated as an adapted HEMIT implementation rather than an exact reproduction of the original model.

### 4.4 H&E–ST Representation Learning

#### (a) Model architecture

The H&E–ST branch was designed to learn a transcriptome-associated modality-aligned representation, termed UR2, from paired H&E images and DeepSpot-derived pseudo-ST targets. The branch contained independent modality-specific encoders for H&E and pseudo-ST. The H&E encoder received a 3 × 256 × 256 image and consisted of four stride-2 downsampling blocks with channel dimensions of 32, 64, 128, and 256, followed by two residual blocks, producing an H&E-derived UR2 of dimensions 256 × 16 × 16.

The pseudo-ST encoder received the normalized 16 × 16 × 256 expression tensor. The gene dimension was first projected using a 1 × 1 convolution followed by Group Normalization and GELU activation, and the resulting feature map was processed by two residual blocks to generate a pseudo-ST-derived UR2 with the same dimensions of 256 × 16 × 16.

Both representations supported self- and cross-modal reconstruction, yielding four routes: H&E→H&E, H&E→pseudo-ST, pseudo-ST→H&E, and pseudo-ST→pseudo-ST. The H&E decoder used four twofold bilinear upsampling blocks followed by a 3 × 3 output convolution and sigmoid activation. The pseudo-ST decoder consisted of two residual blocks, a 1 × 1 channel projection, and a linear transformation over the gene dimension to recover a 16 × 16 × 256 expression tensor. No output activation was applied to the pseudo-ST decoder because the normalized pseudo-ST targets contained signed continuous values.

#### (b) Training objectives

The H&E–ST branch was jointly optimized using reconstruction and weak representation-alignment objectives. H&E reconstruction used the same combination of pixel-wise L_1_ and SSIM losses defined for the H&E–mIHC branch, whereas both H&E→pseudo-ST and pseudo-ST→pseudo-ST reconstruction used mean squared error.

To weakly align the H&E- and pseudo-ST-derived UR2 representations without enforcing element-wise correspondence, global average pooling was applied over their spatial dimensions followed by mean squared error,

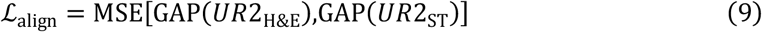

The complete branch objective was

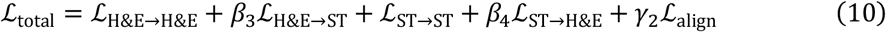

In the final configuration, β_3_=1, β_4_=0.1, and γ_2_=0.05. The reduced weight assigned to pseudo-ST→H&E reconstruction reflected its auxiliary role, whereas H&E→pseudo-ST prediction constituted the principal cross-modal objective.

#### (c) Model training

The H&E–ST branch was trained using the 3717 HEMIT training patches and monitored on the 630 validation patches, while the 945 held-out test patches were excluded from optimization and checkpoint selection. The model was trained for 100 epochs using AdamW with a learning rate of 1×10^-4^, weight decay of 1×10^-5^, and a batch size of 4.

Checkpoint selection was based exclusively on H&E→pseudo-ST reconstruction performance on the validation set. Specifically, the checkpoint with the lowest validation H&E→pseudo-ST mean squared error was retained. The selected model was subsequently used to extract H&E-derived UR2 representations for the downstream UR fusion and biological analysis.

### 4.5 Transfer to Experimentally Measured Spatial Transcriptomics

#### (a) Visium HD data preparation and spatial splitting

Transfer to experimentally measured spatial transcriptomics was evaluated using the 10x Genomics “Visium HD Spatial Gene Expression Library, Human Colorectal Cancer (FFPE)” dataset [22]. Expression profiles were analyzed at the 16 μm aggregated-bin resolution [23]. Counts were normalized as log(1+CP10K), where library size was calculated from all gene-expression features for each bin. Of the 256 genes used by the CUBE H&E–ST branch, 251 were available in the Visium HD dataset; missing genes were excluded using a fixed gene mask.

The tissue section was partitioned into 571 paired H&E–ST patches, each covering 256 μm × 256 μm and containing a 16 × 16 grid of Visium HD bins. H&E patches were normalized toward the HEMIT staining domain using slide-level Macenko normalization followed by global Lab L* calibration [38]. The section was then divided into contiguous spatial stripes along the column axis, with one-block guard bands separating the training, validation, and test regions. A fixed subset of 100 training patches containing at least 128 valid bins was selected for adaptation, while 119 validation patches and 111 held-out test patches were retained for model selection and final evaluation, respectively. The pretrained H&E→UR2 encoder was frozen throughout all real-ST experiments.

#### (b) Zero-shot evaluation and DeepSpot comparison

Zero-shot CUBE evaluation used the pretrained H&E→UR2 encoder and the original pseudo-ST-trained decoder without any optimization on experimentally measured ST. Predictions were converted back from the pseudo-ST normalization used during CUBE pretraining and compared directly with the measured Visium HD expression targets.

DeepSpot was evaluated independently on the same 111 held-out test patches using the same normalized H&E input domain, spatial grid, gene ordering, valid-bin mask, and matched-gene set. This ensured that CUBE and DeepSpot were compared using identical measured targets and spatial regions. Evaluation was performed only over valid Visium HD bins and genes available in both the CUBE output space and the measured dataset.

#### (c) Decoder-only adaptation to real ST

To assess whether the H&E-derived UR2 could be recalibrated to experimentally measured expression, the H&E→UR2 encoder was kept fixed and only the UR2→ST decoder was optimized. Real-ST targets were transformed using gene-wise Z-scores calculated exclusively from the selected training patches, and optimization used masked mean squared error over valid spatial bins and available genes.

Three decoder initialization strategies were evaluated. In the pretrained setting, the complete decoder was initialized from the pseudo-ST-trained CUBE checkpoint and subsequently fine-tuned. In the scratch setting, the complete decoder was randomly initialized while retaining the same frozen UR2 encoder. In the reset-head setting, the pretrained decoder trunk was retained, whereas the final 256→256 linear output layer was reinitialized; all decoder parameters were then jointly optimized. Thus, reset-head differed only in initialization and did not restrict training to the final layer.

Decoder adaptation used AdamW with a learning rate of 1×10^-4^, weight decay of 1×10^-5^, batch size 16, and a maximum of 100 epochs. Validation loss was monitored using early stopping with a patience of 15 epochs and a minimum improvement threshold of 10^-4^.

#### (d) Evaluation metrics and spatial visualization

Real-ST prediction was quantified using gene-wise Pearson correlation, gene-wise Spearman correlation, and bin-wise Pearson correlation on the 111 held-out test patches. Gene-level summary metrics were calculated for genes detected in at least 50 valid test bins; 247 genes met this criterion. All quantitative comparisons used the same 23,710 valid test bins.

For spatial visualization, predictions were additionally generated across all 571 patches and reconstructed according to the native Visium HD grid coordinates. Because whole-section maps from adapted decoders included training and validation regions, these visualizations were used only for qualitative assessment; all reported quantitative performance was derived exclusively from the fixed held-out test region.

### 4.6 UR Fusion and Concept Prediction

#### (a) Model architecture

The UR fusion module integrated the H&E-derived UR1 and UR2 representations extracted from the pretrained H&E–mIHC and H&E–ST branches. For each H&E patch, UR1 had dimensions of 256 × 32 × 32 and UR2 had dimensions of 256 × 16 × 16. The two representations were flattened into 1024 and 256 spatial tokens, respectively, each with an embedding dimension of 256. Independent learnable positional embeddings and summary tokens were added to the two token sequences.

To incorporate the global spatial location of each patch, its normalized two-dimensional coordinate was encoded using fixed Fourier features constructed from sine and cosine functions at 64 logarithmically spaced frequencies [39]. The resulting 256-dimensional coordinate embedding was linearly projected and added to both UR1 and UR2 token sequences.

Each sequence was first processed by an independent self-attention block followed by a feed-forward network. Bidirectional cross-attention was then performed, with UR1 attending to UR2 and UR2 attending to UR1, followed by an additional feed-forward block on each branch [40]. All attention modules used a 256-dimensional embedding space, eight attention heads, pre-LayerNorm residual connections, a feed-forward dimension of 512, and dropout of 0.1 [41,42].

The enhanced summary token from each branch was extracted and concatenated to form a 512-dimensional fused representation. This representation was passed through a concept-prediction head consisting of a 512→256 linear layer, GELU activation, dropout, and a final 256→3 linear layer followed by sigmoid activation, yielding continuous predictions for the DAPI-, CD3-, and panCK-derived concept scores.

#### (b) Training objective

The fusion model was trained exclusively using concept prediction loss. Let 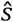 and s denote the predicted and target three-dimensional concept-score vectors, respectively. The objective was

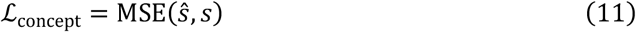

No reconstruction or additional representation-alignment losses were applied during fusion-model training.

#### (c) Model training

UR1 and UR2 features were precomputed using the selected H&E–mIHC and H&E–ST checkpoints and remained fixed during fusion-model training. The fusion module was trained on the 3717 HEMIT training patches and monitored on the 630 validation patches, while the 945 held-out test patches were excluded from model optimization and checkpoint selection.

Training was performed for 80 epochs using AdamW with a learning rate of 1×10^-4^, weight decay of 1×10^-5^, and a batch size of 4. The checkpoint with the lowest validation concept mean squared error was retained for subsequent evaluation and downstream representation analysis.

### 4.7 Representation Ablation and Tissue-Stratified Concept Evaluation

#### (a) Single-representation ablation models

To assess the contribution of UR1 and UR2 to concept prediction, two single-representation ablation models were trained independently rather than generated by masking one branch of the fused model. The UR1-only model received the fixed H&E-derived UR1 representation, whereas the UR2-only model received the fixed H&E-derived UR2 representation. Both models additionally used the same normalized global coordinate information as the fusion model.

Each ablation retained the corresponding positional embedding, summary token, Fourier coordinate embedding, and 256-dimensional attention representation. Because only one representation stream was present, cross-attention was omitted and replaced by successive self-attention and feed-forward blocks before concept prediction. The resulting summary token was passed through the same 256→256→3 concept-prediction head with sigmoid activation. Both ablation models were trained independently using the same concept MSE objective and train/validation split as the full fusion model, with checkpoint selection based on minimum validation MSE.

#### (b) Concept-prediction evaluation

Fusion, UR1-only, and UR2-only models were evaluated using identical concept targets and data partitions. Overall performance was quantified by mean squared error across the three mIHC-derived concept scores, together with marker-specific Pearson correlations for DAPI, CD3, and panCK. The total Pearson correlation was calculated by concatenating the predicted DAPI, CD3, and panCK concept scores across all evaluated patches and computing a single Pearson correlation against the correspondingly concatenated target scores. A constant baseline defined by the mean concept-score vector of the training set was additionally evaluated using the same protocol. All model selection was performed on the validation set. The held-out test set was used only after checkpoint selection to assess generalization of the independently trained fusion and ablation models.

#### (c) Tissue-stratified sensitivity analysis

Because concept-prediction performance was sensitive to the proportion of tissue contained within individual H&E patches, an additional tissue-stratified sensitivity analysis was performed on the held-out test set. Tissue fraction was estimated directly from H&E intensity without histological segmentation. Pixels for which all three RGB channels were at least 245 were defined as near-white background, and tissue fraction was calculated as

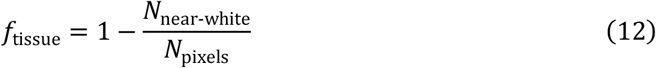

Evaluation was repeated after retaining patches with tissue fractions of at least 1%, 5%, 10%, 25%, 30%, 50%, and 75%, in addition to the full test set. At each threshold, Fusion, UR1-only, and UR2-only were compared on exactly the same retained patches using the same concept-prediction metrics. To summarize the relative advantage of multimodal fusion, the MSE difference was defined as

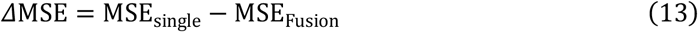

such that positive values indicate lower prediction error for Fusion. The complete threshold series was retained as a sensitivity analysis rather than selecting a single tissue-content cutoff post hoc.

### 4.8 Latent-Space Visualization and Spatial Clustering

To visualize the geometry of the fused CUBE representation independently of explicit global-coordinate information, all 5292 HEMIT patches were re-encoded using a single fixed coordinate equal to the mean normalized coordinate across the complete dataset. The resulting 512-dimensional fused representations were reduced to 50 principal components and embedded into two dimensions using UMAP with 30 neighbors, a minimum distance of 0.25, Euclidean distance, and random seed 2026 [24,25].

The same PCA-reduced representations were additionally subjected to K-means clustering with K=8 to obtain a finer-grained exploratory partition of the latent space for spatial visualization [26]. Cluster assignments were mapped back to their original HEMIT parent-region grids using the patch-level spatial indices encoded in the sample names, yielding patch-level spatial cluster maps rather than pixel-level tissue segmentations.

### 4.9 Immune–Epithelial Spatial Enrichment Analysis

#### (a) Immune–epithelial spatial enrichment score

To quantify local spatial association between immune and epithelial signals, an immune–epithelial (IE) enrichment score was derived from the original 1024 × 1024 mIHC images. CD3 and panCK channels were independently binarized using Otsu thresholding [18]. For each CD3-positive pixel, the local fraction of panCK-positive pixels was calculated within a circular neighborhood of radius 20 μm, corresponding to 80 pixels at 0.25 μm per pixel. These local fractions were averaged across all CD3-positive pixels and normalized by the global panCK-positive fraction of the same patch. The IE score was defined as

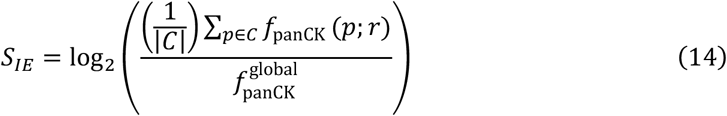

where C denotes the set of CD3-positive pixels and f_panCK_(p;r) is the panCK-positive fraction within radius r=20 μm around pixel p. Positive values indicate local panCK enrichment around CD3-positive regions relative to the patch-wide panCK abundance, whereas negative values indicate relative depletion. Patches lacking either CD3- or panCK-positive pixels, or otherwise yielding an undefined enrichment value, were excluded.

#### (b) Representation probing

The extent to which IE enrichment was encoded by different CUBE representations was assessed using independently trained MLP probes. Frozen UR1 and UR2 representations were spatially averaged to obtain 256-dimensional feature vectors, whereas the 512-dimensional fixed-coordinate fused representation described in Section 4.8 was used directly. Each representation was mapped to the IE score using the same MLP architecture with hidden dimensions of 128 and 64, GELU activations, dropout of 0.1, and a scalar output.

Probes were trained for 80 epochs using AdamW, mean squared error loss, a learning rate of 1×10^-4^, weight decay of 1×10^-5^, and batch size 128. The checkpoint with the lowest validation MSE was retained, and held-out test performance was evaluated using Pearson correlation, Spearman correlation, and R^2^.

#### (c) Control baselines

Two additional controls were evaluated using the same MLP architecture and training protocol. The Concat baseline concatenated spatially averaged UR1 and UR2 into a 512-dimensional vector without attention-based fusion. The Concepts baseline used only the three DAPI, CD3, and panCK Otsu-positive pixel fractions as input. These controls were used to distinguish information captured by the fused representation from that obtainable through simple feature concatenation or marker abundance alone.

### 4.10 External ECM-Associated Transcriptomic Program Analysis

#### (a) ECM-associated transcriptomic program definition

An extracellular matrix (ECM)-associated transcriptomic program score was calculated from experimentally measured Visium HD expression using the KEGG ECM–receptor interaction pathway (hsa04512) [27]. Seventeen pathway genes were present in both the CUBE gene space and the measured Visium HD dataset: COL1A1, COL1A2, COL4A1, COL4A2, COL6A1, COL6A2, COL6A3, COMP, FN1, HSPG2, ITGA11, ITGA5, SPP1, THBS1, THBS2, TNC, and VWF.

For each patch, log(1+CP10K) expression was first averaged across valid 16 μm aggregated-bins for each gene. Gene-wise means and standard deviations were then calculated exclusively from the 294 training-region patches and used to Z-score expression values across all patches. The ECM program score for patch ii was defined as the mean standardized expression across the 17 genes,

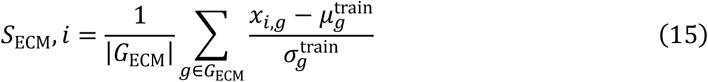

where x_i,g_ denotes the patch-level expression of gene g, and μ_g_ ^train^ and σ_g_ ^train^ were derived from the training region only.

#### (b) Representation probing

The ability of different CUBE representations to predict the ECM program score was assessed using the same MLP probing framework described in Section 4.9. Spatially averaged UR1 and UR2 provided 256-dimensional inputs, whereas the 512-dimensional fused representation was generated using a fixed global coordinate to remove direct dependence on absolute patch location.

Independent probes were trained for UR1, UR2, and Fusion using the spatially separated Visium HD split comprising 294 training patches, 119 validation patches, and 111 held-out test patches; the 47 intervening gap patches were excluded from probe training and quantitative evaluation. Model selection was based on minimum validation MSE.

#### (c) Held-out evaluation and spatial visualization

Probe performance was evaluated on the 111 held-out test patches using Pearson correlation, Spearman correlation, and R^2^. For spatial visualization, measured and Fusion-predicted ECM scores were additionally reconstructed across all 571 Visium HD patches according to their native spatial coordinates, with the held-out test region indicated separately. Whole-section maps were used for qualitative visualization, whereas quantitative correlations and scatter plots were calculated exclusively from the held-out test region. Test-set Pearson correlations for UR1, UR2, and Fusion were additionally summarized for direct representation-level comparison.

### 4.11 Implementation and Statistical Analysis

#### (a) Implementation and computational environment

CUBE was implemented in Python 3.12.2 using PyTorch 2.10.0 with CUDA 12.8. Model training and inference were performed on NVIDIA L40 GPUs. Numerical and statistical analysis were conducted using NumPy, SciPy, scikit-learn, and pandas, with Matplotlib and UMAP-learn used for visualization and latent-space analysis where applicable. Random-number generators in Python, NumPy, and PyTorch were initialized with a seed of 2026 where stochastic procedures were used.

#### (b) Statistical analysis

Pearson correlation was used to quantify linear agreement, Spearman correlation to assess rank-based association, and R^2^ to quantify explained variance where applicable. Mean squared error was used as the principal optimization and model-selection criterion for continuous concept and representation-probe prediction tasks. Unless otherwise stated, quantitative performance was calculated on the predefined held-out test samples or spatial test regions described above. Paired comparisons were performed using matched patches or genes evaluated by both methods.

## Author’s Contribution

Z.G. and H.C. contributed to the conception and design of the project. H.C. supervised the project and provided resources. Z.G. developed and implemented CUBE, conducted all computational experiments and analyses, curated the data, prepared the figures, and interpreted the results. Z.G. wrote the original manuscript, and H.C. reviewed and edited the manuscript.

## Acknowledgments

Z.G. thanks Zheyuan Dong and Ronghao Cao for their encouragement and informal discussions during the development of this work.

## Data Availability Statement

The source data supporting the findings of this study are publicly available on Zenodo at https://doi.org/10.5281/zenodo.22713021. The archive contains figure-associated source data, processed results, and supporting metadata underlying the analyses reported in this study. Raw third-party datasets are not redistributed and remain available from their original providers under the corresponding access and licensing terms. The HEMIT dataset used in this study is publicly available through Mendeley Data at https://data.mendeley.com/datasets/3gx53zm49d/1 and the official HEMIT repository at https://github.com/BianChang/HEMIT-DATASET. The experimentally measured colorectal cancer Visium HD dataset used for transfer evaluation and downstream transcriptomic analysis is the 10x Genomics “Visium HD Spatial Gene Expression Library, Human Colorectal Cancer (FFPE)” dataset, available at https://www.10xgenomics.com/datasets/visium-hd-cytassist-gene-expression-libraries-of-human-crc-v4.

## Code Availability

The source code for CUBE, including data preprocessing, model training, evaluation, representation analysis, and downstream analyses, is publicly available at https://github.com/Zhiyan-Ge/CUBE. Pretrained CUBE model weights, benchmark checkpoints, and downstream probe checkpoints associated with this study are archived on Zenodo at https://doi.org/10.5281/zenodo.22713201. The Zenodo metadata record is publicly accessible, whereas access to the archived weight files is restricted because redistribution and reuse of the weights may be subject to applicable third-party model licensing terms.

## Use of Generative Artificial Intelligence

During the preparation of this study and manuscript, OpenAI ChatGPT was used to assist with English-language translation and language editing of author-prepared text, as well as code development and organization. OpenAI ChatGPT Images 2.0 was used to generate schematic elements for Figure 1 and to generate the Table-of-Contents (ToC) graphic, which were subsequently assembled and edited by the authors. All AI-assisted code, scientific analyses, interpretations, and manuscript content were reviewed, verified, and approved by the authors. No experimental or biological image data were generated or modified using generative artificial intelligence.

